# Macroevolutionary bursts and constraints generate a rainbow in a clade of tropical birds

**DOI:** 10.1101/489419

**Authors:** Jon T. Merwin, Glenn F. Seeholzer, Brian Tilston Smith

**Affiliations:** Department of Ornithology, American Museum of Natural History, Central Park West at 79th Street, New York, NY 10024, USA; Department of Ecology, Evolution and Environmental Biology, Columbia University, New York, NY 10027, USA

**Keywords:** macroevolution, bird, color, phylogeny, model adequacy, lorikeet, mosaic evolution

## Abstract

**Background:** Bird plumage exhibits a diversity of colors that serve functional roles ranging from signaling to camouflage and thermoregulation. However, birds must maintain a balance between evolving colorful signals to attract mates, minimizing conspicuousness to predators, and optimizing adaptation to climate conditions. Examining plumage color macroevolution provides a framework for understanding this dynamic interplay over phylogenetic scales. Plumage evolution due to a single overarching process, such as selection, may generate the same macroevolutionary pattern of color variation across all body regions. In contrast, independent processes may partition plumage into sections and produce region-specific patterns. To test these alternative scenarios, we collected color data from museum specimens of an ornate clade of birds, the Australasian lorikeets, using visible-light and UV-light photography, and comparative methods. We predicted that the diversification of homologous feather regions, i.e., patches, known to be involved in sexual signaling (e.g., face) would be less constrained than patches on the back and wings, where new color states may come at the cost of crypsis. Because environmental adaptation may drive evolution towards or away from color states, we tested whether climate more strongly covaried with plumage regions under greater or weaker macroevolutionary constraint.

**Results:** We found that alternative macroevolutionary models and varying rates best describe color evolution, a pattern consistent with our prediction that different plumage regions evolved in response to independent processes. Modeling plumage regions independently, in functional groups, and all together showed that patches with similar macroevolutionary models clustered together into distinct regions (e.g., head, wing, belly), which suggests that plumage does not evolve as a single trait in this group. Wing patches, which were conserved on a macroevolutionary scale, covaried with climate more strongly than plumage regions (e.g., head), which diversified in a burst.

**Conclusions:** Overall, our results support the hypothesis that the extraordinary color diversity in the lorikeets was generated by a mosaic of evolutionary processes acting on plumage region subsets. Partitioning of plumage regions in different parts of the body provides a mechanism that allows birds to evolve bright colors for signaling and remain hidden from predators or adapt to local climatic conditions.

## Introduction

Animals and plants express a dazzling range of colors. Color has a direct impact on fitness through signaling [1–5], camouflage [2–4], and thermoregulation [6–8], and is a key signal of adaptive diversification and constraint. For birds in particular, plumage color plays a key role in many aspects of their diverse life histories, with notable evolutionary consequences. The major factors which drive the evolution of plumage color are climatic adaptation, crypsis, and sexual selection [4, 9]. Sexual selection is often invoked particularly to explain the evolution of extreme ornamentation and colorfulness seen in various groups of birds [4,10,11]. Examining the macroevolutionary trends of plumage within brightly colored clades provides a framework for understanding how natural and sexual selection interact over phylogenetic scales [9,12,13].

Typical avian clades with ornamental traits show extreme sexual dimorphism, in which males exhibit exaggerated features in form and color as compared to females, which generally have mottled brown or gray cryptic coloration [11]. In contrast, the brightly colored parrots (Order: Psittaciformes) are among the gaudiest of birds but are predominantly monomorphic [14]. As opposed to colorful dichromatic groups such as the birds of paradise, there is little direct evidence that any one factor such as strong sexual selection drives parrot plumage evolution, although some work suggests that assortative mating has driven color evolution in Burrowing Parrots [15, 16]. Although colorful feathers may appear maladaptively conspicuous, parrot feather pigments have been linked to antibacterial resistance, solar radiation protection, and anti-predator defense [17]. While the characteristic bright green displayed by most parrots is decidedly cryptic against a leafy background [18, 19], it is unclear whether sexual selection or drift alone have generated and partitioned the rest of the color gamut in Psittaciformes. Phylogenetic relationships among all parrots are reasonably well known [20], yet few subclades have the dense taxon-sampling necessary for detailed comparative analysis. The one exception is the the brush-tongued parrots, or lories and lorikeets (Tribe: Loriini; hereafter lorikeets) [21]. Lorikeets have radiated into over 100 taxa across the Australasian region [14] since their origin in the mid-Miocene [22]. In comparison to other groups of parrots, lorikeets are species-rich given their age [22]. Their rapid diversification was likely driven by allopatric speciation as they dispersed across Australasia and may be linked to the evolution of their specialized nectarivorous diet [23]. Lorikeets have evolved an extraordinary spectrum of plumage colors which range from vibrant ultraviolet blue to deep crimson and black. These colors are organized in discrete plumage regions or patches which in turn vary in size, color, and placement among taxa yet are nonetheless easily defined and compared across species.

The macroevolutionary patterns that underlie the radiation of these color patches in lorikeets can provide context into how diverse coloration evolves. As with many complex multivariate traits (e.g., [24–26], we expect that mosaic evolution, wherein subsets of traits evolve independently of others, underlies bird plumage color diversification. Different color metrics (e.g., hue vs. brightness) may be under independent selective pressures to balance a tradeoff between eye-catching ornamentation and cryptic background matching [12, 18]. For example, in the Eclectus parrot (*Eclectus roratus*) the males have bright green plumage for camouflage against predators while foraging and moving between mates, and the females have bright red and purple coloration to advertise nest-sites [18]. In the predominantly monomorphic lorikeet taxa, however, color variation appears to be partitioned along a dorso-ventral axis with the head, breast and abdominal feather regions being more variable than the wings and back. If the level of color variation is linked to whether the plumage region was subject to natural or sexual selection, or drift then distinct macroevolutionary patterns should be observable among patches. While assessing the relative fit of different macroevolutionary models cannot ascribe process, comparing their likelihoods will determine whether the distribution of color among taxa and plumage regions is consistent with a model of mosaic evolution.

In this study we quantified and modeled color evolution in the lorikeets to test whether plumage in this group is evolving as a mosaic or as a simple trait evolving under a similar evolutionary rate on all body regions. To produce color data, we imaged museum specimens, extracted spectral reflectance from plumage regions, and summarized color hue, disparity, and volume. We tested whether dorso-ventral partitioning of plumage regions can explain color evolution in lorikeets by fitting alternative evolutionary models using comparative phylogenetic methods. We predict that the relatively low color variation of dorsal plumage regions has been structured by natural selection for crypsis, and should be best explained by a model where there is a cost to evolving to new color states. In contrast, the variable face and ventral patches are likely involved in conspecific signaling and therefore evolutionary change in these patches would be expected to carry less cost, i.e., they would radiate under lower constraint. Alternatively, if plumage has evolved due to a single overarching process, selection or drift might dictate the evolutionary trajectory of color variation for all patches simultaneously. Under this type of scenario, we would expect all patches to be explainable by the same model. Because color is often correlated with environmental conditions [27–30], we interpreted our modeling selection results in the context of the relationship plumage color and climatic variables. Characterizing the veritable rainbow of colors in the lorikeets and testing alternative scenarios that could give rise to this variation will help clarify whether discrete macroevolutionary patterns have partitioned color diversification or whether a single model will best explain the color variation in all color patches.

## Results

### Macroevolutionary model selection

We found that independent patterns or rates have indeed generated color variation in the lorikeets, but our results were more nuanced than our proposed alternative scenarios. One extra level of complexity was that best fit models for individual patches varied among principal component (PC) axes. The first principal component (PC1, representing 52% of variance) of color primarily represented brightness, meanwhile the second (PC2, 27%) and third principal components (PC3, 13%) represented hue in the blue-to-red and UV-to-Green axes, respectively (Supplementary Figure S3). In PC1 (achromatic variation or brightness), Delta and Brownian Motion models were best fit to the dorsal patches of the wings, back, and crown. The breast and face patches however were best fit to Ornstein-Uhlenbeck (OU) models. Hue principle components showed the opposite pattern. PC2 (blue-to-red chromatic variation) for the forehead, crown, and occiput was best fit by Brownian Motion models with lambda (λ) values of one, indicating a rate of evolution equal to the expected signal under a random walk along the phylogeny. Face, breast, and tail evolution was best supported by a Delta (δ) model. All other patches for PC2 were best supported by an OU model. The best-fit model for most patches was selected with high relative support by sample-size corrected Akaike Information Criterion, (ΔAIC_C_ >4) except for crown, forehead and occiput (Δ AIC_C_ < 2; Supplementary Figure S1). For PC2, wing, wrist, rump, and breast were best fit by an OU model. The best-fit model of PC2 for lower abdominal patches, lateral neck, and tail was Brownian Motion, while an OU model explained half of the wing, wrist, eyeline, and lower breast color. All other patches, which were clustered around the abdomen, head, and face, were best modeled by a late-burst Delta model. We found that most best-fit models were a good absolute fit to the patch color data (Figure 3F). Undescribed rate heterogeneity in the back, crissum, and secondaries (PC2) caused model-adequacy to fail for these patches. When we assessed model adequacy by comparing statistics estimated from empirical and simulated trait values, we used a four-statistic threshold for determining absolute fit, but many patches would have passed a five- or six-statistic threshold (Supplementary Table S1). Most best-fit models were robust to simulation tests and our absolute adequacy filter in arbutus (Supplementary Table S2) [31]. Of the six calculated model adequacy statistics, C*_var_*, the coefficient of variation of the paired differences between the estimated node and tip values, most frequently deviated from empirical values (Supplementary Table S2).

**Figure 1:**
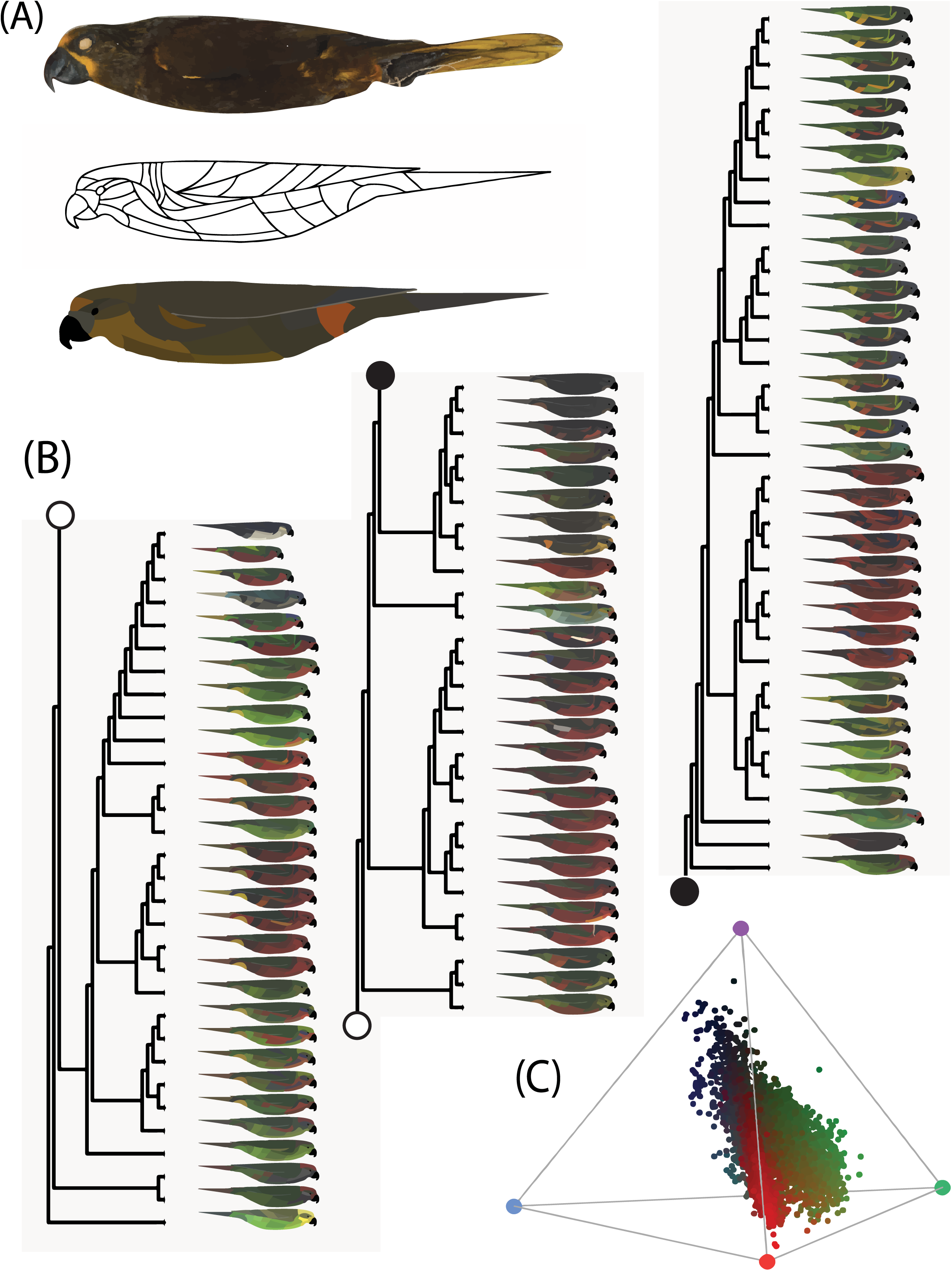
Quantifying and plotting plumage color on a phylogeny of lories and lorikeets. (A) An image of a museum specimen of *Chalcopsitta duivenbodei* (top), a blank patchmap showing the 35 plumage regions measured from images of museum specimens (middle), and the corresponding patchmap for this exemplar taxon (bottom). (B) Patchmaps of all taxa (*n* = 98) plotted on a phylogeny. The tree was split into three sections and the connecting portions are indicated with corresponding filled or empty points. (C) The tetrahedral color space of the Loriini, which contains four vertices for the four measured reflectance wavelengths: UV (purple, top), short (blue, left), medium (green/yellow, right), and long (orange/red, center). Each point represents one of the 35 color patch measurements for each taxon. The color space was centered slightly towards the longwave (red) vertex of the tetrahedral color space. While the distribution of colors in the color space skews towards the longwave part of the spectrum, it was most variant in the UV spectrum and also exhibits wide variance in the medium-wave spectrum. Colors represent the RGB colors which were mapped onto the real-color patchmaps.

**Figure 2:**
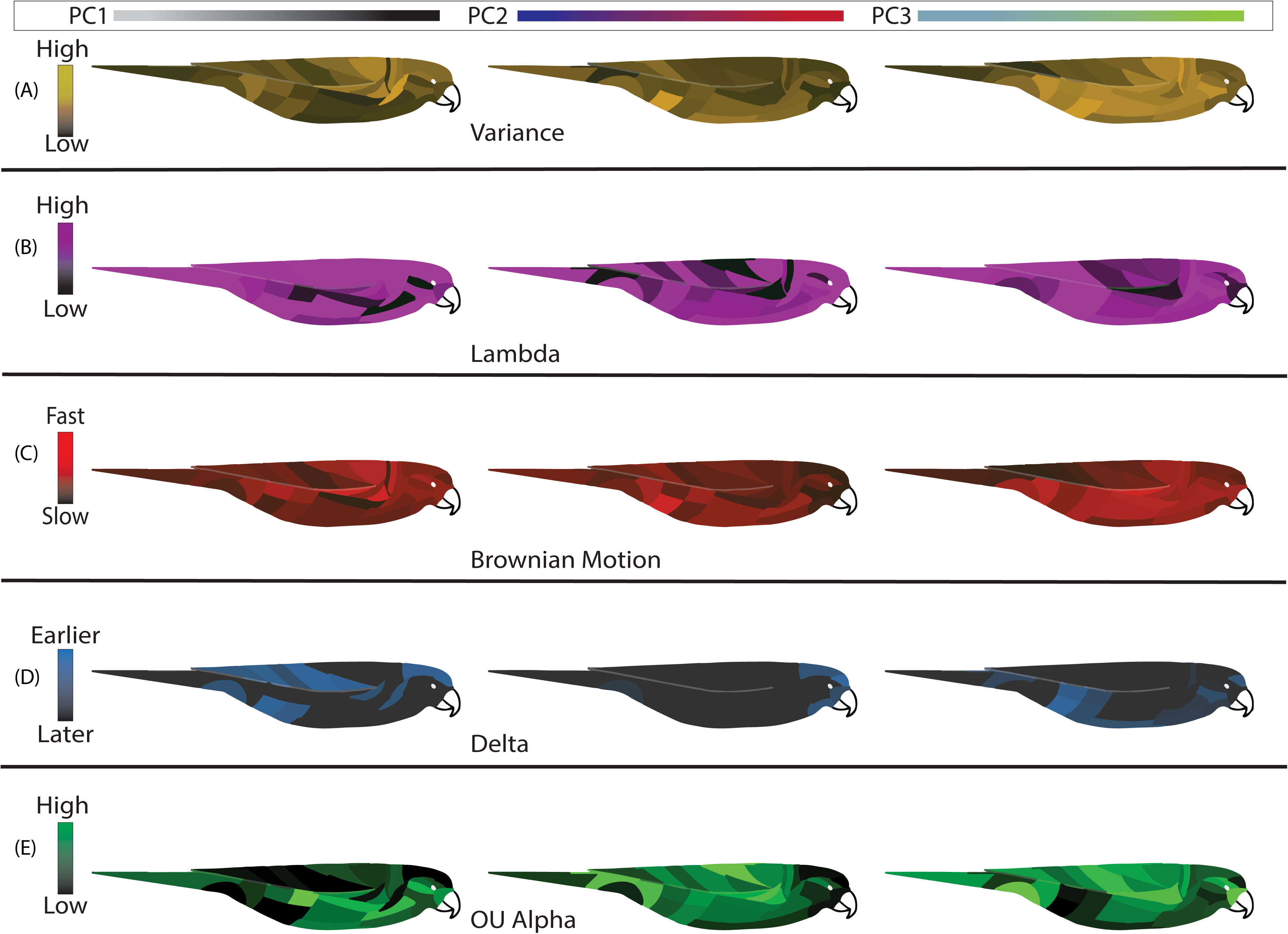
Patchmaps show within and among patch variability in PC values, model parameters, and best-fit models. PC scale bars at top show axes of color variance encompassed by each PC. Each patch in a patchmap was colored according to values for principal component variance (A), the modeled parameters lambda (B), Brownian Motion rate (C), delta (D) and OU alpha (E), and the best-fit model, after model adequacy (F). The left and right patchmaps within each panel represent PC1 and PC2, respectively. From top to bottom, the darker patches are less variable across taxa (A), have less phylogenetic signal (B), are evolving slower (C), diversified closer to the tips of the tree (D), or were relatively more constrained (E). See Supplementary Table S2 for a full listing of model-fit parameters.

**Figure 3:**
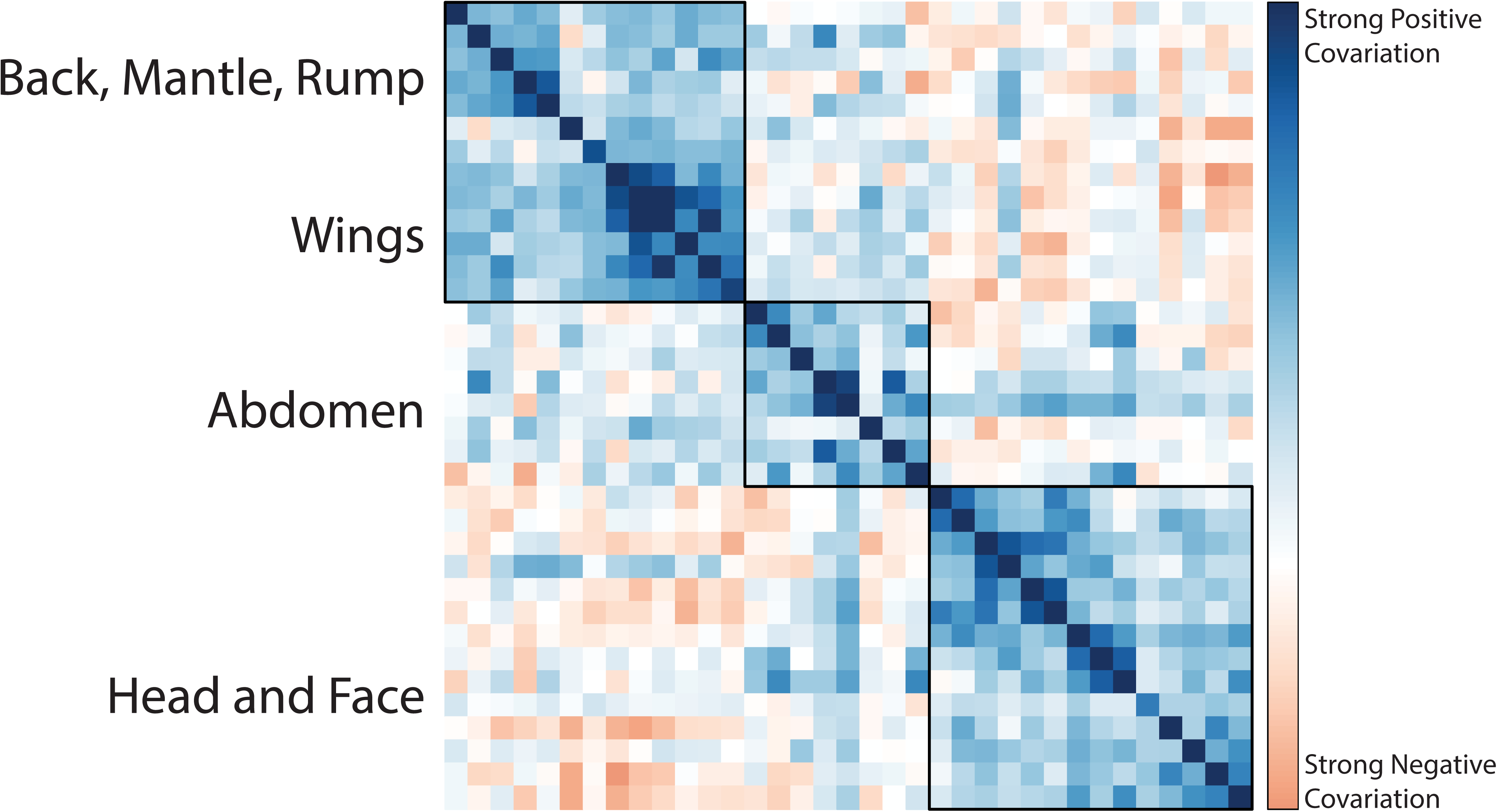
Phylogenetic variance-covariance matrix across all 35 patches shows that patch color in discrete morphological regions covary. Darker blue colors represent stronger positive covariance while darker red colors represent stronger negative covariance. Boxes represent hierarchical clusters, estimated using the built-in hclust method in corrplot.

Multi-trait, non-independent model fitting on all 35 patches showed that the highest-likelihood multi-trait model was an OU model, suggesting that all patches were under constraint. However, the variance-covariance matrix of this model fit showed hierarchical clustering of covariant patches on the head, abdomen, and wing (Figure 3). When we tested alternative scenarios of trait grouping, we found that three separately evolving modules on the face, breast, and wing were the maximum likelihood scenario (Figure 4D). Multi-trait model fitting on only these correlated patch subsets indicated that head and breast patch variation was best explained by a Delta model, while for wing and abdominal patches an OU model was recovered. Visualizations of the variance-covariance matrix of patches showed that patches which were best-fit by similar models during the individual patch analysis also covaried under a single all-patch model (Figure 3).

**Figure 4:**
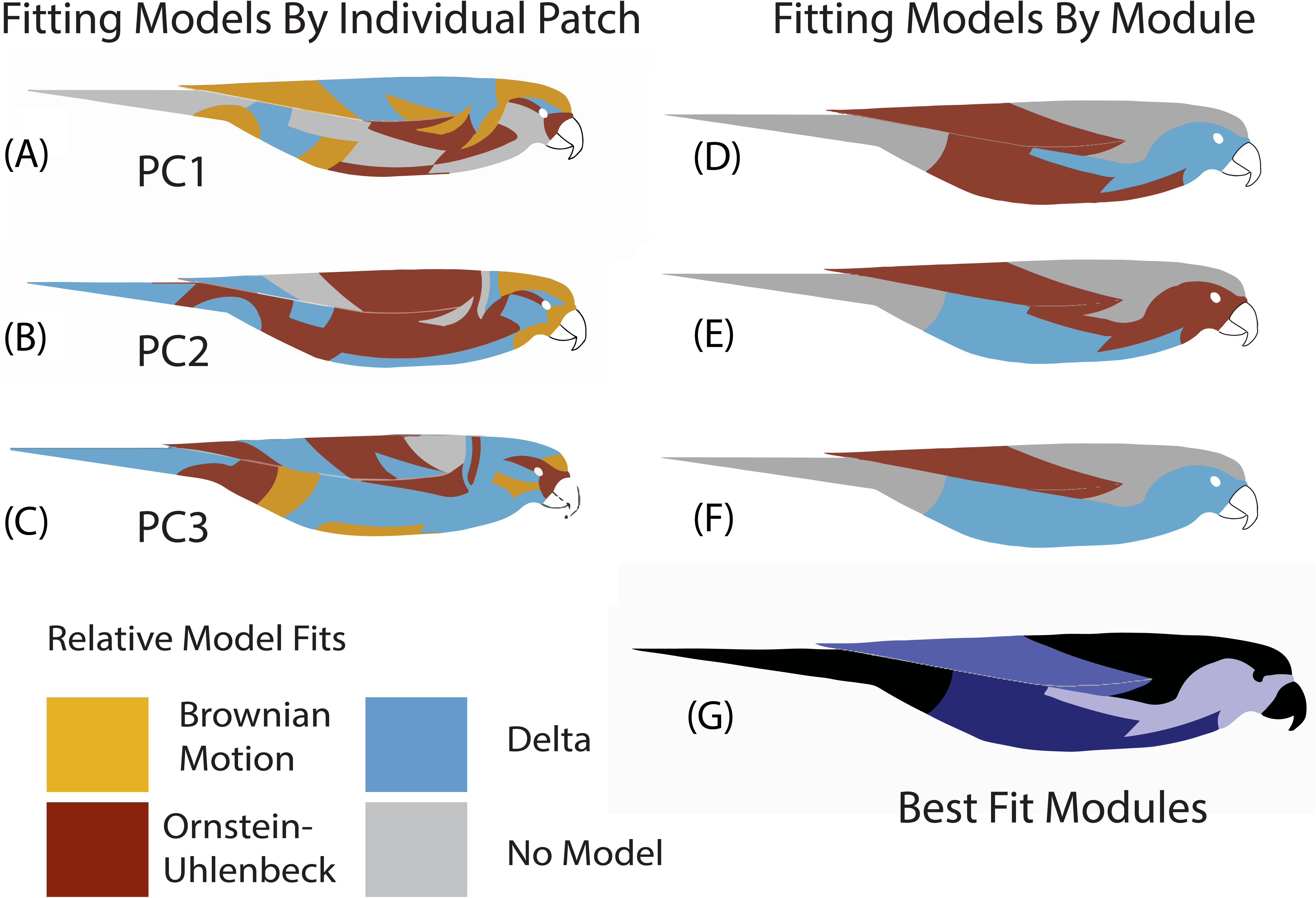
Relative model fits show a mosaic of best-fit models across patches for PC1-PC3 (A-C), and that most patches were a good absolute fit to the data. Different colors represent different evolutionary models and only patches with good absolute fits were plotted. For the Models fit by module (D-F), models were fit for PC1 (D), PC (E), and PC3 (F) of color for patch groups outlined in the maximum likelihood scenario (G). Note that no patches were best fit by White Noise models.

Overall, values were high, suggesting that color is a strong signal of phylogenetic relatedness. An examination of the parameters estimated from all tested models shows how phylogenetic signal (the extent to which phylogeny explains trait variation) varied among patches and hue and brightness. For all principal components and for most patches, fitted values were at the upper bound of the metric, indicating that phylogenetic signal was equal to the expected signal under Brownian Motion (Figure 2B, Supplementary Table S2). For color PC1 (brightness), the malar region, and patches along the side (side breast) had the lowest phylogenetic signal (Figure 2B). In contrast, the back, wrist, and crissal patches exhibited the lowest phylogenetic signal for color PC2 and PC3. The fastest rate of evolution of PC1 was detected in the back, wrist, and abdominal patches (Figure 2C). All patches fit a Delta model with δ > 1, indicating that every patch followed a late-burst pattern of evolution (Figure 2D). For PC2, many model δ values were at the default maximum, 3. For PC3, δ was 3 for the wings, body, crown and crissum, but lower on the tail, back and side-throat. OU alpha (α) values showed a similar pattern. High α values, which represent stronger pull towards an estimated optimum, were fit to lower abdomen patches, wings, and wrists. The areas under weakest constraint (low α) were the breast, face, and head patches.

### Ancestral Reconstruction

Patch colors on the face and abdomen change from node to node, while similar wing colors (mainly green) are distributed across the tree and are generally conserved between nodes (Figure 5B, Supplementary Figure S2). We visualized this pattern using continuous color mapping of single patches versus wing chord length (Supplementary Figure S2). We found repeated evolution of patch colors across distantly related genera and high color divergence between closely related genera. However, morphometric traits such as wing chord length exhibited less heterogeneous evolutionary rates (Supplementary Figure S2), largely reflecting that the taxa within genera have similar body sizes.

**Figure 5:**
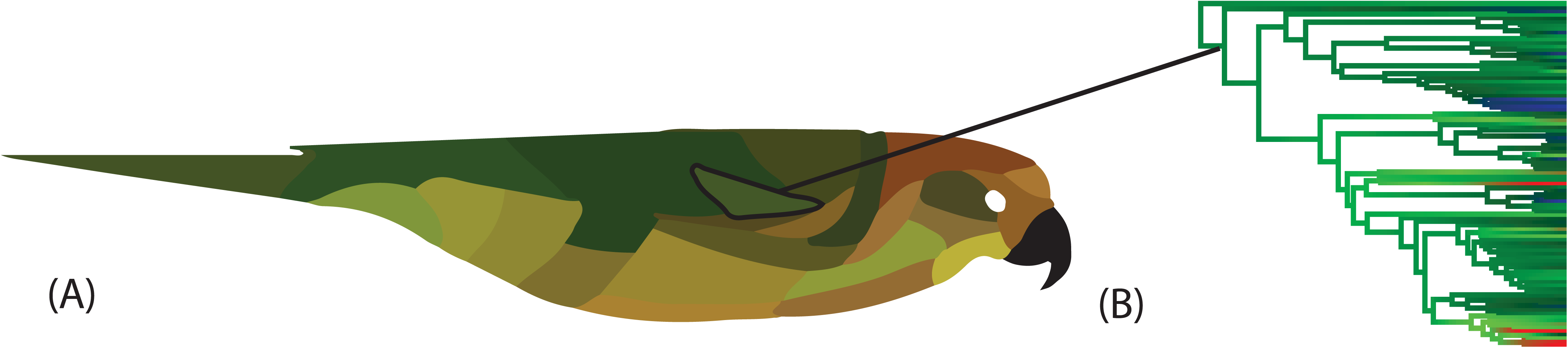
Visualization of ancestral reconstruction of all patches at the basal node to all lorikeets. Note the lighter-colored underside countershading to the dark green dorsal wings and back, which are conserved across deep nodes and found in most extant lorikeets. (B) represents an example of an ancestral reconstruction of a single patch (the lesser coverts) with an arrow pointing to the node from which ancestral states were extracted for each patch. While the colors in B were approximated based on a single PC axis, the ancestral colors reported in (A) were calculated based on RGB and UV reflectance simultaneously.

We constructed a patchmap of mean ancestral states of all patches using the anc.recon method in phylopars [32]. The resulting ancestral lorikeet had dark green wings, a lighter green-yellow torso, a reddish crown and forehead, and blue cheek patches, closely resembling aspects of both *Trichoglossus chlorolepidotus* and *Charmosyna rubronotata* (Figure 5A). Ancestral patchmaps plotted using the maximum or minimum of the 95% CI were qualitatively similar to those made with the mean ancestral color value and were not plotted. To further visualize how color evolved across the tree, an animation of ancestral state patchmaps from the root of the lorikeets to the basal node of *Lorius lory lory* (Electronic Supplement 1) is provided as an exemplar.

### Color and Climate

Lorikeets occupied 33.5% of the colors predicted to be perceivable by tetrachromatic birds. The average color volume per taxon was 0.00513, which represents a relative volume of around 2.37% (median 2.07%) of the total avian visual space. Individual taxon color volumes ranged between 0.04% to 11.7% of avian visual space. The average largest pairwise distance between two patches for one bird, or average hue disparity, was 0.912 (median: 0.962).

A phylogenetic generalized least-squares (PGLS) analysis modeling the relationship between climate and color found nuanced patterns that showed that color was correlated with precipitation, temperature, and elevation, but the nature of these relationships varied between modeling all patches at once or patches in distinct patch regions. We report multivariate models that were selected using AIC values, which were assessed as we sequentially removed insignificant model variables (with highest p-values) until the AIC of the new model was not significantly statistically different than the previous model, in which case the previous model was selected (ΔAIC < 2). All models had the same number of response variables (*n* = 75). Full variable loadings for each climate principal component are available in the supplementary material (Supplementary Figure S6).

When we modeled all patches as a response variable and principal components of climate as predictors, brighter feathers (lower color PC1) were correlated with warmer (Temperature PC2) and drier (Precipitation PC2) environments (R^2^ = 0.12; *p* < 0.05; Table 2). Overall lorikeets were greener (higher color PC3) in areas with higher precipitation seasonality (R^2^ = 0.08; *p* < 0.05; Table 2). Climate did not explain a significant portion of blue-to-red variation (color PC2). Wing color was more correlated with climate than face or abdomen when we correlated small groups of patches with climate (as opposed to modeling the color of all patches at once). On the wings, greener color was associated with higher seasonality and higher temperatures (R^2^ = 0.17; *p* < 0.001). On the abdomen, we found that darker plumage was associated with lower temperatures, higher precipitation seasonality, and low elevation (R^2^ = 0.09; *p* < 0.03). Wing patch hue PC1 (R^2^ < 0.01; *p* = 0.37), abdomen hue (R^2^ =0.04; *p* = 0.10), and face patches overall (R^2^ = 0.001; *p* = 0.3) were poorly predicted by climate or elevation.

## Discussion

The evolution of the exceptional color variation in lorikeets was best explained by independent patterns or rates acting on different plumage regions and axes, namely brightness and hue. Overall, both independent patch and correlated patch subset analysis showed that while some plumage regions were drawn towards optimum values over time, others diversified along the phylogeny in bursts, suggesting that different plumage regions are subject to alternative evolutionary regimes. As is the case with many traits that characterize color [4], the plumage of lorikeets was only partially explained by climatic variation, but those patches that covaried with temperature and/or precipitation were conserved across the phylogeny. In contrast, those patches that evolved in bursts and were not associated with climatic variation may be evolving in response to sexual selection, social selection, or drift. Collectively, our results suggest that at a phylogenetic scale, lorikeet plumage color has evolved in correlated regions, a pattern consistent with the idea that natural and sexual selection independently acted on components of a multivariate phenotype.

### Functional underpinnings of mosaic evolution

Our results suggest that plumage evolution has been partitioned between the back and the front (dorso-ventral axis) and between the face and the rest of the body in the lorikeets, indicating that the patterns that govern plumage evolution vary with regard to location on the body of an organism. Selected best-fit models clustered in independent units on the face, breast, and wing; these regions are readily interpretable based on our functional knowledge of plumage color biology. For instance, the crown, forehead, and lower abdomen were best supported by a model of Brownian Motion, which may be because these regions are under processes such as sexual or social selection. One taxon in our dataset, *Trichoglossus haematodus* is known to flare and preen their bright crown and forehead feathers during courtship, but any specific role color plays in this display is not well known [33]. An OU model fit to hue for most wing and body patches is consistent with either a constraint on evolution to new hue states for climatic adaptation, or cryptic background matching. In the forest canopy, green body and wing color may serve the purpose of camouflage against predation [6, 34], while brighter plumage colors may serve as signals, as observed in the reversed sexually dichromatic parrot *Eclectus roratus* [18]. Highly variable and colorful regions, like the face, breast, and tail, were best explained by a Delta model both in the individual-patch and module model fitting. Our inferred δ parameters were greater than one, which indicates color variance within these patches evolved towards the tips of the tree. Although this pattern can be interpreted as evidence for character displacement [35, 36], the majority of taxa within clades are currently allopatric [14], so recent color evolution was presumably unaffected by interactions with other lorikeet taxa. Instead, rapid bursts of evolution across many color patches likely reflects the commonly observed pattern of rapid color evolution at the tips of phylogenies, which may indicate that these patches may function as signals to conspecifics or may be under sexual selection [13, 37]. In those lorikeets that do exhibit sexual dichromatism (e.g., some taxa in *Charmosyna*), the face patches are the regions that vary in color [14].

The difference in evolutionary dynamic that we observed between lorikeet face and wing patches may be driven by divergent selective forces. Within the Loriini and across Psittaciformes, green wings are a common phenotype [14, 37], as 90% of parrots have green patches and 85% are primarily green [38]. The fact that wing patches were best explained by an OU model may indicate there is a selective cost to evolving away from green. Species with green wings and backs are predicted to have increased camouflage in trees against aerial and terrestrial predators [18]. While we found a correlation between climatic factors and color on the wings and the abdomen, this pattern did not hold for face patches. In contrast to monochromatic birds, which may be under strong selection for uniform plumage color (such as the snow-colored winter plumage of Rock Ptarmigans; [39], lorikeet faces may be colorful, in part, because their color variation is not constrained by natural selection. Highly hue-variable regions, like the breast and face, were not explained well by an OU model, suggesting that there has been no “optimum” value for the hue of these patches across the radiation of lorikeets. Therefore, these small, variable facial patches and bright breast patterns present across the Loriini may be important signals to conspecifics, while monochrome green dorsal feathers may provide cover from predators against green canopy backgrounds.

Overall, we found that the direction and magnitude of color-climate relationships differed between principal components of color and between plumage regions. Discrete body regions showed divergent association patterns between hue and climate. Across all patches, birds were brighter in seasonal, dry areas, and darker in wet areas, supporting Gloger’s Rule [30, 40]. In lorikeets, brightness and hue may be subject to different forces, a pattern which has been chiefly observed in less chromatically variable birds. Overall, the strongest relationship that we found was between wing color greenness and temperature, precipitation, and elevation. Our results suggest that birds at higher elevations and in warmer temperatures had greener wings. While wing color was most correlated with climate, abdomen and face patches showed a less pronounced or no pattern, suggesting that ornamental and cryptic coloration in lorikeets are balanced along the dorso-ventral axis.

### Model adequacy

Overall, all our best-fit models had good absolute fit. Prior work based on non-color traits found that relative models fit to subclades within a family-level phylogeny (the Furnariidae) had good absolute fit, but these same models had poor absolute fit when applied at the family scale [31, 41]. In our dataset, simulated values of one statistic (C*_var_*) frequently deviated from empirical values because of unaccounted-for rate variation in our best-fit, constant rate model. Even at relatively shallow phylogenetic scales, body size and plumage color exhibit rate heterogeneity [25,41,42]. Accounting for rate shifts by testing the Delta model was critical for accurately characterizing the evolution of highly variable regions, which may be rapidly shifting between several discrete states or diversifying due to sexual selection.

### Independent or correlated patches

The developmental architecture that underlies potential concerted evolution among feather regions remains unknown for most birds [43, 44]. We found that there were three clusters of correlated patches that correspond to adjacent sections on the wing, breast, and face (Supplementary Figure S3). We found that these clusters were correlated when hierarchically clustered in a phylogenetic variance-covariance matrix (Figure 5) and when analyzed in a phylogenetically-naive likelihood framework against alternative clustering hypotheses (Figure 4c). These regions may be developmentally linked, under similar selective regimes, or the result of differential regulation of separate genes across patches or patch regions [69]. Regulatory controls on feather color may work at patch-level, feather tract-level, or whole bird-level scales [43–45], and understanding how these pathways are connected will elucidate how complex plumage colors and patterns evolve. For example, most lorikeets have all-green wings with black-tipped primaries, and our ancestral reconstruction analysis suggests that the ancestor to all lories had green wings, but some *Eos* taxa have evolved red wings with black barring and UV coloration on some wing patches, demonstrating a clear interplay between region- and patch-level pigment and structural color regulation. In the sister taxon to lorikeets, *Melopsittacus undulatus*, a single base-pair change expresses tryptophan, blocking expression of yellow pigment, changing the mostly-green wild-type to a pale-blue across all patches [44]. A similar simple molecular change may explain the evolution of the two brilliant blue taxa in the Loriini; *Vini ultramarina* and *V. peruviana*, or the evolution of red-colored lorikeets in the genera *Eos*, *Pseudeos*, and *Trichoglossus* [38].

### Colorful groups have recurring colors

When individual clades radiate across a high percentage of the available color space, then the repeated evolution of similar colors may be a common feature [9, 46]. For example, the robust-bodied and short-tailed lorikeets in *Lorius* and the distantly related, slender and small, long-tailed lorikeets in *Charmosyna* both have red bodies and green wings. Ancestral states inferred from ancestral character estimation in phylopars [32], while subject to a high degree of uncertainty (Supplemental Figure S2), suggest that green wings may have been historically conserved across this radiation and red bodies have originated multiple times. Despite lorikeets being exceptionally colorful, their radiation was not characterized by constant gain of new colors, but rather repeated evolution of similar colors across the phylogeny. Novel color evolution in birds is modulated by interactions between genes, gene expression patterns, structures, and existing metabolic pathways [43,44,47]. Biochemical constraints likely played a role in this plumage convergence because parrot feather color is controlled via regulatory pathways as opposed to dietary pigmentation [48]. Certain trait shifts, such as loss of ancestral yellow/green pigments and gains of red, are common in lorikeets and across parrots [38]. In carotenoid-based color systems such as in the songbird genus *Icterus*, a relatively small number of color states rapidly oscillate, leading to convergence in carotenoid and melanosome-based colors [49, 50]. A similar process may be occurring in lorikeets despite the chemically unique pigmentation found in Psittaciformes. Regardless of mechanism, architectural constraints on plumage color or morphological traits may produce similar looking but distantly-related taxa.

### Challenges in studying plumage color

Quantifying color from museum specimens presented numerous challenges. Using museum specimens instead of hand-painted plates from field guides was preferable to us because skins exhibit UV reflectance, and the three-dimensional variation of the specimen can be captured. However, the variable preparation of museum specimens may expand or obscure certain feather patches. Therefore, we relied on subjective judgement and consultation of multiple skins, plates, and photographs when creating and implementing our patch sampling ontology. Patch outlines were drawn by hand to account for preparation style. One possible solution for patch delineation could be through random sampling of patch location [51]. The potential error in our approach pertains mostly to patch delineation, not the overall color volume of the entire bird. Despite our concerns about the subjectivity in identifying the location of patches on specimens, much of the potential error was likely minimized because of the overall morphological similarity across our focal clade as well as the fact that we performed most elements of our analysis on correlated patch groups. Additionally, patchmaps and field guide plates were qualitatively similar. In studies that sample across much deeper phylogenetic scales, identifying and sampling homologous patches will be a much more complicated task. Machine learning approaches, possibly guided by evo-devo data on feather color and pattern regulation [45], may lead to more objective patch-specific analyses. Delineating high-contrast boundaries would enable patch geometry and boundaries to be objectively quantified [45, 47] and provide a clearer means of interpreting patch colors in the context of sexual or social signaling.

## Conclusion

We found that alternative macroevolutionary models clustered in three groups on the face, abdomen, and wings best explained the exceptional colour variance in the lorikeets. Such mosaic evolution is consistent with the view that separate selective and stochastic processes help shape different plumage regions and have enabled lorikeets to evolve extreme colours despite the selective costs of conspicuous colouration. Demonstrating that mosaic evolution operates in birds and other animals will clarify how extreme phenotypic diversification occurred under variable evolutionary pressures.

## Materials and Methods

### Specimen imaging, color extraction, and visualization

To quantify color, we photographed the lateral, ventral, and dorsal sides of one male museum skin for 98 taxa deposited at the American Museum of Natural History (Supplementary Table S3). This sampling represents 92% of the described diversity in Loriini, all described genera, and all taxa for which phylogenomic data exists. Specimens were photographed using a Nikon D70s with the UV filter removed and a Novoflex 35mm lens. All specimens were lit using four Natural LightingNaturesSunlite 30-W full spectrum fluorescent bulbs (5500K, 93 CRI) attached to arms mounted to a metal copy stand. Using baader spectrum filters affixed to a metal slider, specimens were photographed in both “normal” Red/Green/Blue (RGB) color as well as in the UV spectrum [22, 52].

We demarcated 35 homologous plumage patches on the images produced for each specimen to quantify the variation among taxa based on examination of specimens, plates, and plumage topography maps (Figure 1A; Supplementary Figure S1). Using the multispectral imaging package (MSPEC) in ImageJ [53] we linearized images in DCRAW and normalized images to five gray standards placed alongside each bird and extracted RGB and UV reflectance for each patch. Linearization and normalization control for light balance, maximize contrast, and standardize photographs to known reflectance values, enabling the extraction of objective color data [53]. Measurements were collected using a bluetit visual model in the MSPEC program, which transformed the data from the camera-specific (Nikon D70s) color space into an objective color space and then into a tetrachromatic avian visual model supplied with MSPEC [53]. Full details of the color-space transformation are available in Troscianko and Stevens (2015). Data was plotted in tetrahedral color space using the R v. 3.4.3 [54] package pavo v. 1.3.1 [55]. Using pavo, we extracted summary statistics (volume, relative volume, hue angle, and hue angle variance) of color spaces at varying phylogenetic scales within the Loriini and generated relative hue variables which were scaled to 1 (Table 1). We also measured wing-chord and tarsus length as proxies for body size.

We visualized color data and model output using a 2D schematic of an outline of a generic lorikeet, hereafter referred to as a “patchmap.” (Figure 1A; [54, 56]) We wrote a custom R script to automatically color our patchmaps with raw reflectance data. Specifically, the script input raw blue, green, and red reflectance data into the RGB method in the R package grDevices version 3.4.3 to generate hex colors for each patch for each taxon [54, 56]. Images were plotted as tip labels on a phylogeny representing all of our sampled taxa [21] (Figure 1C) using ggtree v. 1.10.4 in R [55, 57]. Patchmap image sizes were scaled to represent relative taxa sizes measured from museum skin wing lengths.

### Modeling color evolution across the Loriini tree

We modeled patch color across the phylogeny to visualize phylogenetic signal across patches, and to compare how much particular patches evolved from ancestral states. We predicted that patches linked to crypsis (e.g., wing) would maintain a similar color across the tree. In contrast, patches that are presumably involved in mate signaling (e.g., face and breast) would show greater disparity across the tree. First, we converted the non-ultrametric tree for the Lorinni from Smith et al. (2018) into a time-calibrated tree using the program treePL [58]. To date the tree, we used a secondary calibration from [22] by specifying the age for the node separating the Loriini from their sister taxon *Melopsittacus undulatus* to 11-17 million years ago (Mya). This time-calibrated tree was used in all downstream analyses.

### Ancestral character estimation of all patches

Using the anc.recon method in phylopars [63] we performed a multivariate estimation of mean ancestral states using the four raw reflectance variables and visualized these ancestral states on patchmaps to see which specific colors have been conserved over time. Using multivariate ancestral character estimation along multiple axes was necessary because color is a fundamentally multivariate trait and ancestral estimates of single color parameters, such as brightness or a single principal component of hue, cannot account for covariation and independence between different color spectra. We layered images of ancestral states at sequential nodes into animated GIFs to visualize color conservation and shifts for an example sequence of nodes from the root to an exemplar tip (Electronic Supplement 2).

### Macroevolutionary model selection and adequacy test

We tested whether plumage color evolution across patches in lorikeets was best explained by multiple or a single macroevolutionary model. Complex traits are often the product of different evolutionary processes; e.g., tetrapod cranial and postcranial skeletal morphology are subject to discrete forces associated with diet and locomotor strategy, respectively [59]. However, in some cases, a single model may best explain the evolution of a complex trait under strong natural selection, such as with cryptic coloration in female passerines [28]. To test these alternative hypotheses, we used a comparative phylogenetic method to select relative and absolute best-fit models for the first three principal components of color. We compared and selected best-fit models for each patch independently, for correlated patches together, and finally for all patches together using AIC_C_ weights. To analyze color variation across all patches we performed a principal components analysis of all 4,620 color measurements with prcomp in R, using the four raw quantum catch variables (UV, short-, medium-, and long-wave) as factors (R Core Team, 2017). This flattens a four-dimensional color-space matrix into PCs that explain brightness (% of total variance) and color-opponent coordinates (Supplementary Figure S3). For each individual patch, we modeled PC1, PC2, and PC3 for each patch with Brownian Motion, OU, Delta, and White Noise across the phylogeny with the fitContinuous method in the geiger package in R [60] (version 2.0.6). From these models, for each patch we extracted the fit parameters Brownian Motion rate, delta rate-change (δ), phylogenetic signal (λ), and OU bounding effect (α) to assess how parameter values varied across patches independent of their best-fit model. To identify the relative best-fit model for each patch, we considered ΔAIC_C_ greater than 2 to be significantly different. These models were used to test the following expectations: colors evolved as a random walk along phylogeny (Brownian Motion), color evolved within selective constraints (OU), color evolved in a random pattern, irrespective of phylogeny (White Noise), or color evolved in late or early burst fashion (Delta).

Though model selection based on AIC_C_ identifies the best relative model, that model may be overfit and unrealistic [31]. To test model fit adequacy, we compared our empirical trait values to simulated trait values using the arbutus package [31] (version 0.1). Based on a fitted model, arbutus creates a unit tree (a tree with uniform branch lengths of 1), simulates posterior distributions, and compares those simulated distributions of six statistics (Supplementary Table S2) to the empirical trait distribution. When simulated values differed from empirical values (two-tailed P-value; alpha = 0.05), the model had poor absolute fit. We then filtered out the models which failed two or more tests [31, 41]. The best-fit model for each patch was plotted on the patchmap. For patches with ΔAIC_C_ scores of < 2 among top models, the model with fewer parameters was selected. For models with identical complexity, Mahalanobis distance to simulated trait means was used as a post-hoc test in order to pick best-fit models [31, 61].

Because treating individual patches separately in tests may lead to model misspecification due to independent analysis of non-independent traits [62], we fit models for multiple patches at once using the R package phylocurve [63]. This analysis allowed us to identify highly correlated groups of patches and estimate the best-fit model for these correlated groups and all patches together. We generated a phylogenetic covariance matrix for all patches in phylocurve, which fits single trait evolution models to high-dimensional phenotypic traits. These covariance matrices were visualized using the corrplot package [64]. From this covariance matrix we identified correlated patches (e.g., patches on the wing), re-ran phylocurve on these patch clusters, for BM, OU, delta, and white noise models, following the same AIC model selection procedure described in the single patch analysis. To identify the global (entire bird) best-fit model, we compared alternate models fit in phylocurve performed on all patches simultaneously.

### Testing for climatic correlates with color

To test if plumage variation covaried with ecogeographical gradients, we examined the relationship between temperature, precipitation, elevation, and patch color. Although some aspects of lorikeet color, as with other ornamental clades (e.g., [12], may not strongly covary with climate gradients, we aimed to test whether this decoupling of climate and color was present when we tested individual patch regions. Overall, we expected to find that regions involved in climatic adaptation or crypsis (e.g., wings) covary with climatic variables more strongly than regions potentially involved in signaling (e.g., face).

We used the extract function in the raster package to extract the median value from each of 19 bioclim variables [68] as well as elevation [65] from the shapefiles representing each taxon’s distribution [66]. Median bioclim and elevation values were calculated from all raster cells within each taxon distribution shapefile due to a paucity of accurate occurrence records for many lorikeets. We then used the PGLS method in the R package caper [67] to test the relationship between each PC axis for three groups of patches (wing, abdomen, and face) as well as all patches at once, using elevation and the first three principal components of temperature and precipitation as predictors while accounting for phylogeny and using maximum likelihood estimates for lambda. All PGLS models had the same sample size (*n* = 75), which was a reduction from our total taxon list because we excluded subspecies with incomplete range data. We selected the best model for each patch group using AIC values, which were assessed as we sequentially removed insignificant model variables (with highest P values) until AIC of the new model was not 2 lower than the previous model AIC (variables with the highest p-value).

Table 1: Color space statistics for all sampled taxa. All statistics were calculated within a tetrahedral color space using relative reflectance for UV, Short, Medium, and Longwave reflectance. The taxon which occupied the greatest volume of color space was *Phigys solitarius* but taxa in *Trichoglossus* comprised a large portion of the 30 taxa with the greatest color volume. The taxa which occupied the smallest amount of color space were *Vini peruviana*, which is mostly blue, and several *Chalcopsitta* taxa which are monochrome black, brown, and dark red.

Table 2: Best fit PGLS models. Overall, wing patches were best predicted by climate, while we found no relationship between face color and biogeographic variables. Models were selected for each patch subset and each color principal component. Coefficients are presented in the order that they are listed under the “predictors” column, with the intercept value as the first coefficient.

## Supporting information

Supplementary Lorius Animation

## Ethics Approval and Consent to Participate

Not Applicable

## Consent for Publication

Not Applicable

## Availability of Data and Materials

The datasets generated and/or analysed during the current study, as well as a list of specimens used in the color quantification are available in the GitHub repository, https://github.com/jtmerwin/LorikeetColor

## Acknowledgements

We would like to thank S. Simpson for measuring skins, W. Mauck, M. Pennell, F. Burbrink, L. Joseph, E. Morrison, R. Maia, J. Troscianko, M. Palmer, D. H. Brainard, K. Provost, L. Moreira, L. Musher, G. Rosen, T. Trombone, and P. Sweet, and the reviewers and editors for improving the manuscript.

## Literature Cited

1. Edler AU, Friedl TWP. Plumage Colouration, Age, Testosterone and Dominance in Male Red Bishops (Euplectes orix): A Laboratory Experiment. Ethology. 2010.

2. Stevens M, Marshall KLA, Troscianko J, Finlay S, Burnand D, Chadwick SL. Revealed by conspicuousness: distractive markings reduce camouflage. Behav Ecol. 2012;24: 213–222.

3. Gluckman T-L, -L. Gluckman T, Cardoso GC. The dual function of barred plumage in birds: camouflage and communication. J Evol Biol. 2010;23: 2501–2506.

4. Hill GE, Hill GE, McGraw KJ. Bird Coloration: Function and evolution. Harvard University Press; 2006.

5. Bennett AT, Cuthill IC, Partridge JC, Lunau K. Ultraviolet plumage colors predict mate preferences in starlings. Proc Natl Acad Sci U S A. 1997;94: 8618–8621.

6. Medina I, Newton E, Kearney MR, Mulder RA, Porter WP, Stuart-Fox D. Reflection of near-infrared light confers thermal protection in birds. Nat Commun. 2018;9: 3610.

7. Beasley BA, Davison Ankney C. The effect of plumage color on the thermoregulatory abilities of Lesser Snow Goose goslings. Can J Zool. 1988;66: 1352–1358.

8. Hill RW, Beaver DL, Veghte JH. Body Surface Temperatures and Thermoregulation in the Black-Capped Chickadee (Parus atricapillus). Physiol Zool. 1980;53: 305–321.

9. Nordén KK, Price TD. Historical Contingency and Developmental Constraints in Avian Coloration. Trends Ecol Evol. 2018;33: 574–576.

10. Saranathan V, Hamilton D, Powell GVN, Kroodsma DE, Prum RO. Genetic evidence supports song learning in the three-wattled bellbird Procnias tricarunculata (Cotingidae). Mol Ecol. 2007;16: 3689–3702.

11. Irestedt M, Jønsson KA, Fjeldså J, Christidis L, Ericson PGP. An unexpectedly long history of sexual selection in birds-of-paradise. BMC Evol Biol. 2009;9: 235.

12. Dunn PO, Armenta JK, Whittingham LA. Natural and sexual selection act on different axes of variation in avian plumage color. Sci Adv. 2015;1: e1400155.

13. Ornelas JF, González C, de Los Monteros AE. Uncorrelated evolution between vocal and plumage coloration traits in the trogons: a comparative study. Journal of Evolutionary Biology. 2009. pp. 471–484.

14. Forshaw JM. Parrots of the World. 2010.

15. Masello JF, Pagnossin ML, Lubjuhn T, Quillfeldt P. Ornamental non-carotenoid red feathers of wild burrowing parrots. Ecological Research. 2004. pp. 421–432.

16. Masello JF, Quillfeldt P. Body size, body condition and ornamental feathers of Burrowing Parrots: variation between years and sexes, assortative mating and influences on breeding success [Internet]. Emu - Austral Ornithology. 2003. pp. 149–161.

17. Burtt EH Jr, Schroeder MR, Smith LA, Sroka JE, McGraw KJ. Colourful parrot feathers resist bacterial degradation. Biol Lett. 2011;7: 214–216.

18. Heinsohn R, Legge S, Endler JA. Extreme reversed sexual dichromatism in a bird without sex role reversal. Science. 2005;309: 617–619.

19. Kane SA, Wang Y, Fang R, Lu Y, Dakin R. How conspicuous are peacock eyespots and other colorful feathers in the eyes of mammalian predators? PLoS One. 2019;14: e0210924.

20. Provost KL, Joseph L, Smith BT. Resolving a phylogenetic hypothesis for parrots: implications from systematics to conservation. Emu - Austral Ornithology. 2018;118: 7–21.

21. Smith BT, Mauck WM, Benz B, Andersen MJ. Uneven missing data skews phylogenomic relationships within the lories and lorikeets. 2018. doi:10.1101/398297

22. Schweizer M, Wright TF, Peñalba JV, Schirtzinger EE, Joseph L. Molecular phylogenetics suggests a New Guinean origin and frequent episodes of founder-event speciation in the nectarivorous lories and lorikeets (Aves: Psittaciformes). Mol Phylogenet Evol. 2015;90: 34–48.

23. Schweizer M, Güntert M, Seehausen O, Leuenberger C, Hertwig ST. Parallel adaptations to nectarivory in parrots, key innovations and the diversification of the Loriinae. Ecol Evol. 2014;4: 2867–2883.

24. Powell BJ, Leal M. Brain evolution across the Puerto Rican anole radiation. Brain Behav Evol. 2012;80: 170–180.

25. Felice RN, Goswami A. Developmental origins of mosaic evolution in the avian cranium. Proc Natl Acad Sci U S A. 2018;115: 555–560.

26. Barton RA, Harvey PH. Mosaic evolution of brain structure in mammals. Nature. 2000. pp. 1055–1058.

27. Delhey K. A review of Gloger’s rule, an ecogeographical rule of colour: definitions, interpretations and evidence. Biol Rev Camb Philos Soc. 2019;94: 1294–1316.

28. Gomez D, Théry. Simultaneous Crypsis and Conspicuousness in Color Patterns: Comparative Analysis of a Neotropical Rainforest Bird Community [Internet]. The American Naturalist. 2007. p. S42.

29. Delhey K, Dale J, Valcu M, Kempenaers B. Reconciling ecogeographical rules: rainfall and temperature predict global colour variation in the largest bird radiation. Ecol Lett. 2019;22: 726– 736.

30. RM Zink JVR. Evolutionary processes and patterns of geographic variation in birds. Current Ornithology. 1986;4: 1–69.

31. Pennell MW, FitzJohn RG, Cornwell WK, Harmon LJ. Model Adequacy and the Macroevolution of Angiosperm Functional Traits. Am Nat. 2015;186: E33–50.

32. Bruggeman J, Heringa J, Brandt BW. PhyloPars: estimation of missing parameter values using phylogeny. Nucleic Acids Res. 2009;37: W179–84.

33. Serpell J. Visual displays and taxonomic affinities in the parrot genus Trichoglossus. Biol J Linn Soc Lond. 1989;36: 193–211.

34. Soma M, Garamszegi LZ. Evolution of patterned plumage as a sexual signal in estrildid finches. Behav Ecol. 2018;29: 676–685.

35. Schluter, Schluter. Ecological Character Displacement in Adaptive Radiation. Am Nat. 2000;156: S4.

36. Hemingson CR, Cowman PF, Hodge JR, Bellwood DR. Colour pattern divergence in reef fish species is rapid and driven by both range overlap and symmetry. Ecol Lett. 2019;22: 190–199.

37. Stoddard MC, Prum RO. Evolution of avian plumage color in a tetrahedral color space: a phylogenetic analysis of new world buntings. Am Nat. 2008;171: 755–776.

38. Nemeśio A. Colour production and evolution in parrots. International Journal of Ornithology. 2001;4: 75–102.

39. Montgomerie R. Dirty ptarmigan: behavioral modification of conspicuous male plumage. Behav Ecol. 2001;12: 429–438.

40. G. CR. Das Prinzip geographischer Rassenkreise und das Problem der Artbildung. Nature. 1929;124: 753–754.

41. Seeholzer GF, Claramunt S, Brumfield RT. Niche evolution and diversification in a Neotropical radiation of birds (Aves: Furnariidae). Evolution. 2017;71: 702–715.

42. Chira AM, Thomas GH. The impact of rate heterogeneity on inference of phylogenetic models of trait evolution. J Evol Biol. 2016;29: 2502–2518.

43. Morrison ES, Badyaev AV. The Landscape of Evolution: Reconciling Structural and Dynamic Properties of Metabolic Networks in Adaptive Diversifications. Integr Comp Biol. 2016;56: 235–246.

44. Cooke TF, Fischer CR, Wu P, Jiang T-X, Xie KT, Kuo J, et al. Genetic Mapping and Biochemical Basis of Yellow Feather Pigmentation in Budgerigars. Cell. 2017;171: 427– 439.e21.

45. Schwochow-Thalmann D. Molecular Identification of Colour Pattern Genes in Birds. 2018.

46. Hofmann CM, Cronin TW, Omland KE. Using Spectral Data to Reconstruct Evolutionary Changes in Coloration: Carotenoid Color Evolution in New World Orioles. Evolution. 2006;60: 1680–1691.

47. Endler JA, Cole GL, Kranz X. Boundary Strength Analysis: Combining colour pattern geometry and coloured patch visual properties for use in predicting behaviour and fitness. 2018.

48. Berg ML, Bennett ATD. The evolution of plumage colouration in parrots: a review. Emu - Austral Ornithology. 2010;110: 10–20.

49. Omland KE, Lanyon SM. Reconstructing plumage evolution in orioles (Icterus): repeated convergence and reversal in patterns. Evolution. 2000;54: 2119–2133.

50. Morrison ES, Badyaev AV. Structure versus time in the evolutionary diversification of avian carotenoid metabolic networks. J Evol Biol. 2018;31: 764–772.

51. Miller ET, Leighton GM, Freeman BG, Lees AC, Ligon RA. Climate, habitat, and geographic range overlap drive plumage evolution [Internet]. bioRxiv. 2018. p. 375261.

52. McKay BD. The use of digital photography in systematics. Biol J Linn Soc Lond. 2013;110: 1– 13.

53. Troscianko J, Stevens M. Image calibration and analysis toolbox - a free software suite for objectively measuring reflectance, colour and pattern. Methods Ecol Evol. 2015;6: 1320–1331.

54. R Core Team. R: A language and Environment for Statistical Computing [Internet]. R Foundation for Statistical Computing, Vienna, Austria; 2017. Available: https://www.R-project.org

55. Maia R, Eliason CM, Bitton P-P, Doucet SM, Shawkey MD. pavo: an R package for the analysis, visualization and organization of spectral data. Tatem A, editor. Methods Ecol Evol. 2013;I. doi:10.1111/2041-210X.12069

56. Dale J, Dey CJ, Delhey K, Kempenaers B, Valcu M. The effects of life history and sexual selection on male and female plumage colouration. Nature. 2015;527: 367–370.

57. Yu G, Smith DK, Zhu H, Guan Y, Lam TT-Y. ggtree: an r package for visualization and annotation of phylogenetic trees with their covariates and other associated data. McInerny G, editor. Methods Ecol Evol. 2017;8: 28–36.

58. Smith SA, O’Meara BC. treePL: divergence time estimation using penalized likelihood for large phylogenies. Bioinformatics. 2012;28: 2689–2690.

59. Esteve-Altava B. In search of morphological modules: a systematic review [Internet]. Biological Reviews. 2017. pp. 1332–1347. doi:10.1111/brv.12284

60. Harmon LJ, Weir JT, Brock CD, Glor RE, Challenger W. GEIGER: investigating evolutionary radiations. Bioinformatics. 2008;24: 129–131.

61. Revell LJ. phytools: an R package for phylogenetic comparative biology (and other things). Methods Ecol Evol. 2011;3: 217–223.

62. Adams DC, Collyer ML. Multivariate Phylogenetic Comparative Methods: Evaluations, Comparisons, and Recommendations. Syst Biol. 2017;67: 14–31.

63. Goolsby EW. Likelihood-Based Parameter Estimation for High-Dimensional Phylogenetic Comparative Models: Overcoming the Limitations of “Distance-Based” Methods. Syst Biol. 2016;65: 852–870.

64. Taiyun Wei VS. corrplot: Visualization of a correlation matrix. 10/2013.

65. Survey USG, U.S. Geological Survey. Shuttle Radar Topography Mission (SRTM) [Internet]. Fact Sheet. 2003. doi:10.3133/fs07103

66. Birdlife International. Bird Species Distribution maps of the World. Cambridge, United Kingdom and Arlington, United States: Birdlife International and NatureServe; 2011.

67. Orme D. The caper package: comparative analysis of phylogenetics and evolution in R. 2018.

68. Fick, S.E. and R.J. Hijmans, 2017. Worldclim 2: New 1-km spatial resolution climate surfaces for global land areas. International Journal of Climatology.

69. Abolins-Abols M, Kornobis E, Ribeca P, Wakamatsu K, Peterson MP, Ketterson E, et al. A role for differential gene regulation in the rapid diversification of melanic plumage coloration in the dark-eyed junco (Junco hyemalis). 2018.

