## Supplemental Figures. for "Macroevolutionary bursts and constraints generate a rainbow in a clade of tropical birds"

**Supplementary Material**

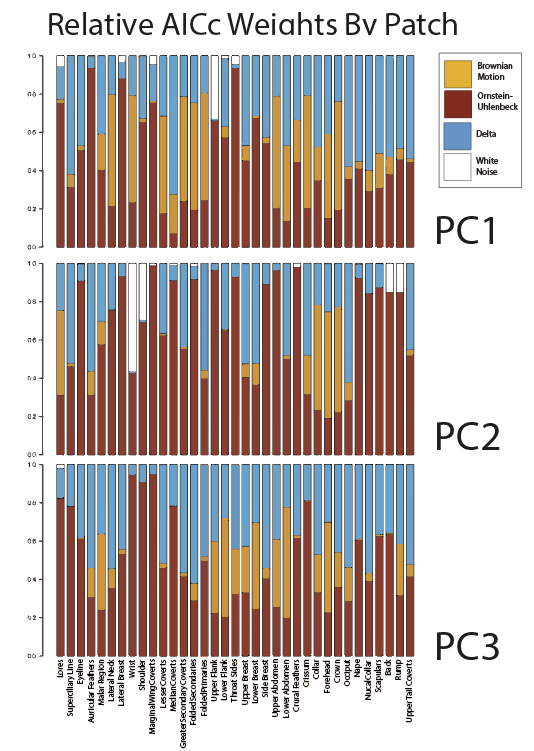

Figure S1: Relative AIC_C_ weights by patch for the by-patch best-fit model selection. Note that in PC1, OU patches were selected with very high relative support. Colors represent the model tested and each bar represents a single patch in a single color PC.

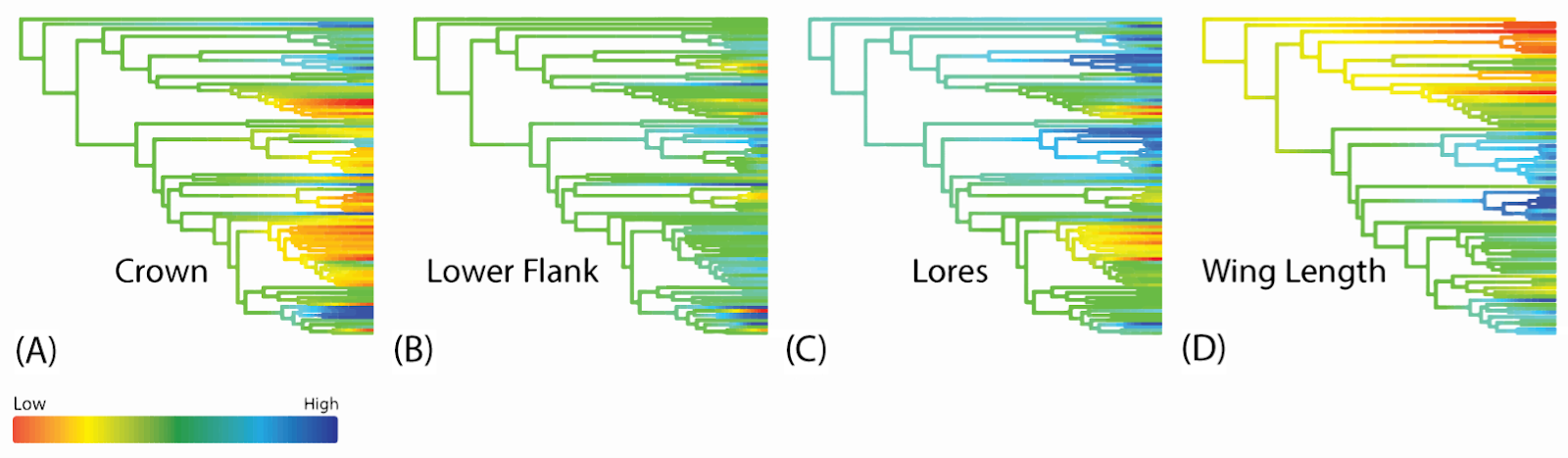

Figure S2: Continuous character mapping shows that color is a more labile trait than body size. Ancestral states were estimated using a Brownian Motion process, and included are exemplar patches and wing length on the phylogeny. The patches correspond to PC1 for the top of the head (A, Crown), lateral view above the legs (B, Lower Flank), area directly next to the bill (C, Lores), and a proxy for body size (D, Wing Length).

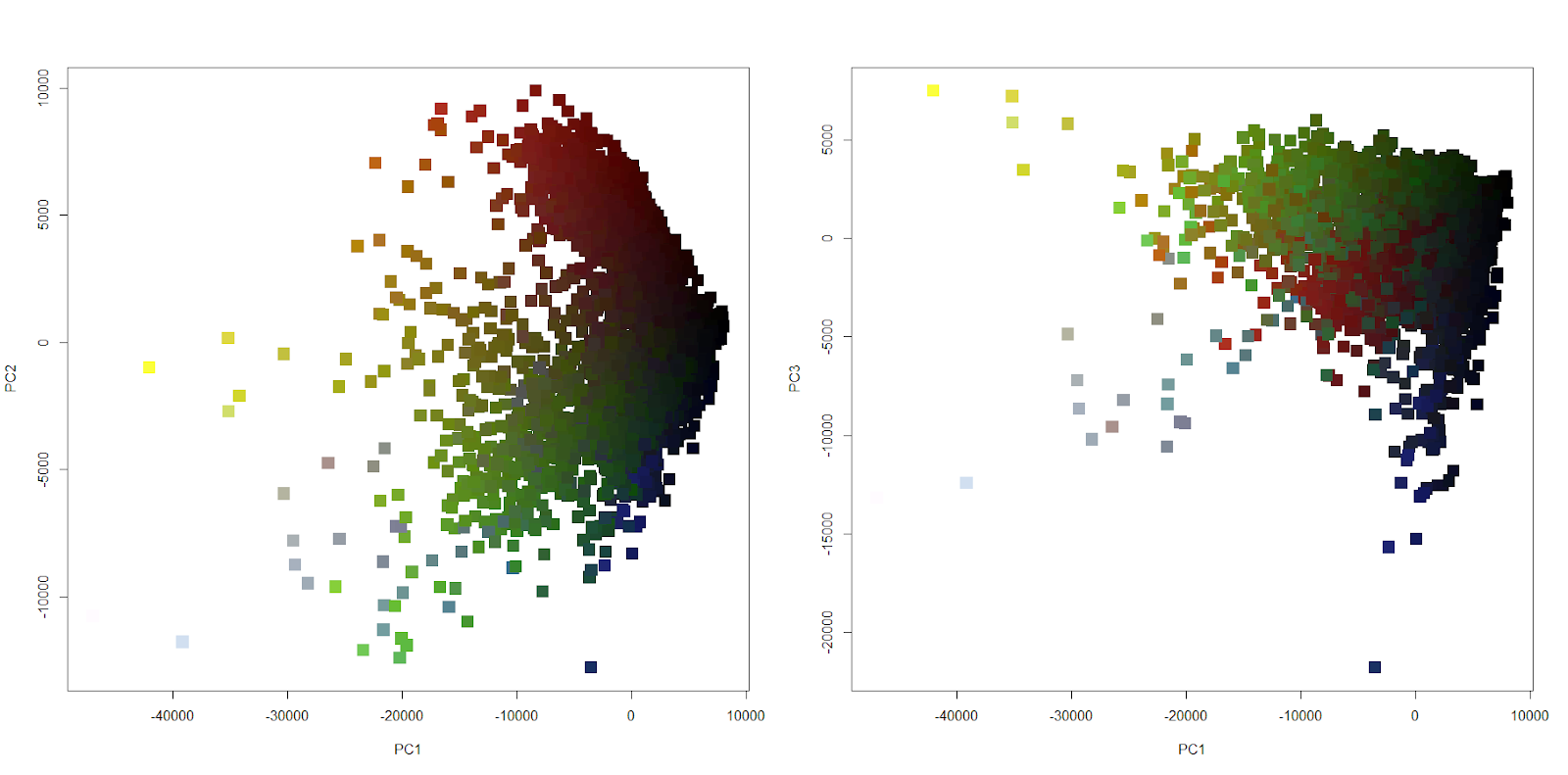

Figure S3: Scatterplot of PC1, PC2, and PC3 of Color. Color on the plots represents real colors that a human would see, as generated by the RGB method in R. This shows that a PCA across all 4,620 color measurements using the four reflectance variables (U, S, M, and L) as factors. Note that PC1 represents achromatic variation (brightness) while PC2 and 3 represent hue variance. PC2 represents variation along the blue-to-red axis while PC3 represents variation along the UV-to-green axis.

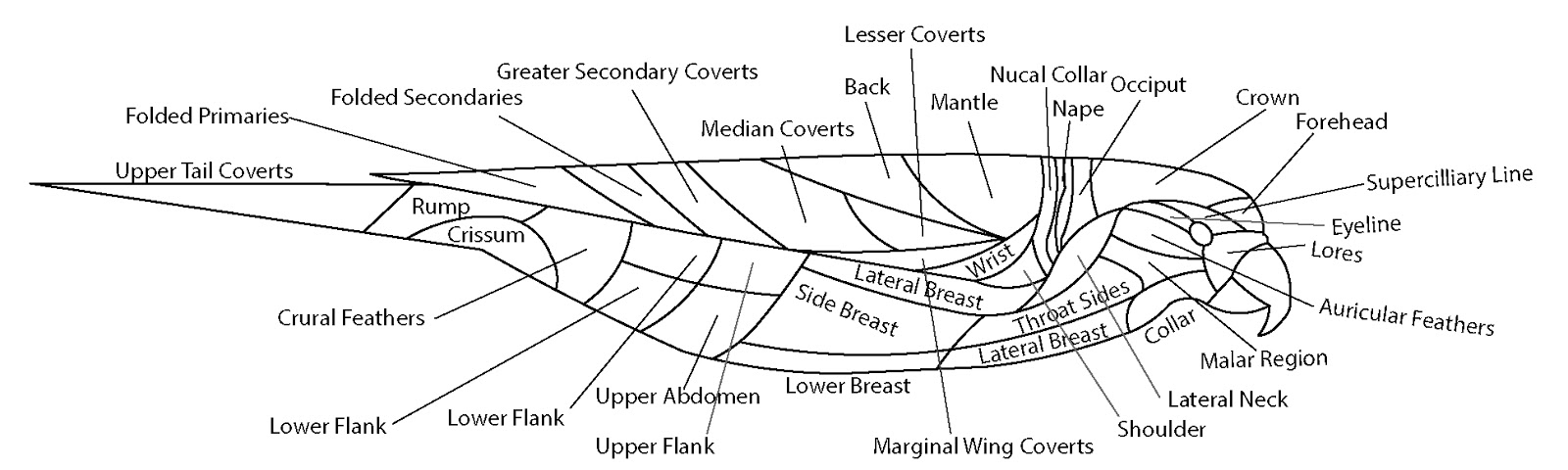

Figure S4: Patchmap showed labeled plumage regions. Color was extracted from these 35 regions on each specimen. The position of patches on specimens was visually approximated and hand delineated on images as per Figure S5.

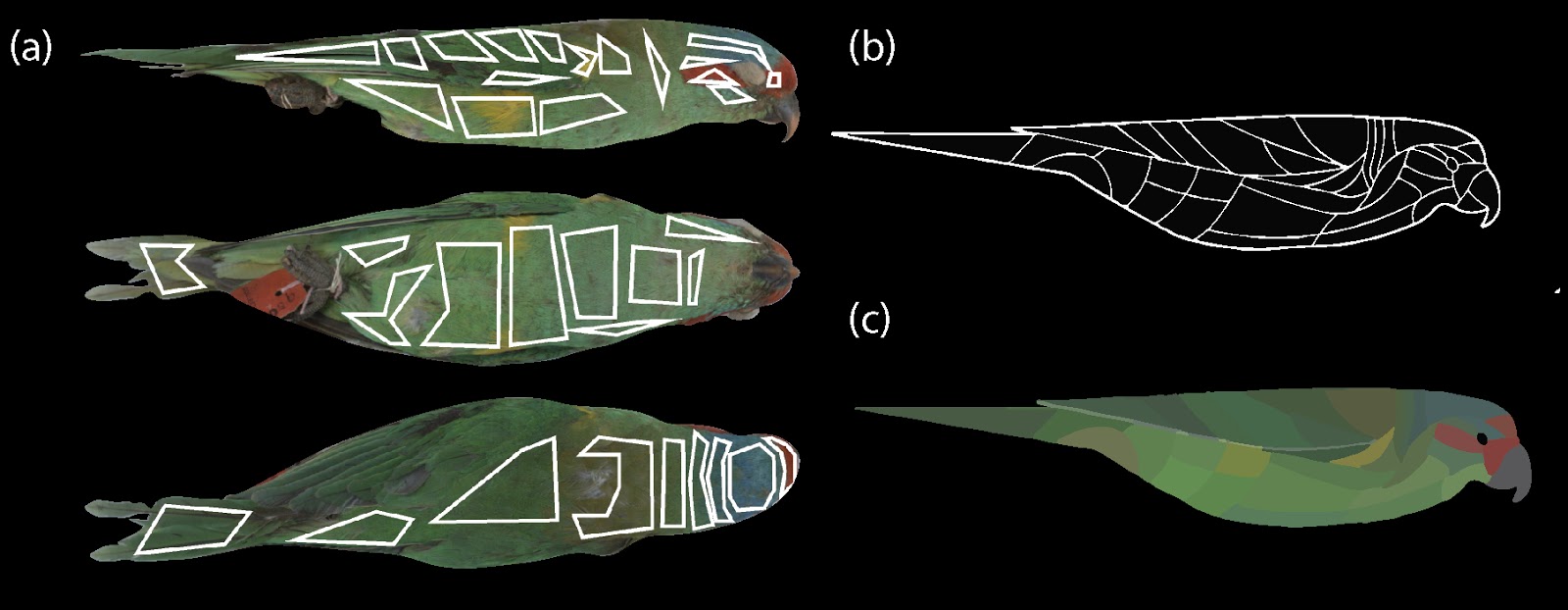

Figure S5: Example of sampling patches from a an example photo (a). Patch sampling scheme was adapted from McKay et al., while examining plates and skins to adequately capture full 3D bird variance. Patches were selected from three views.

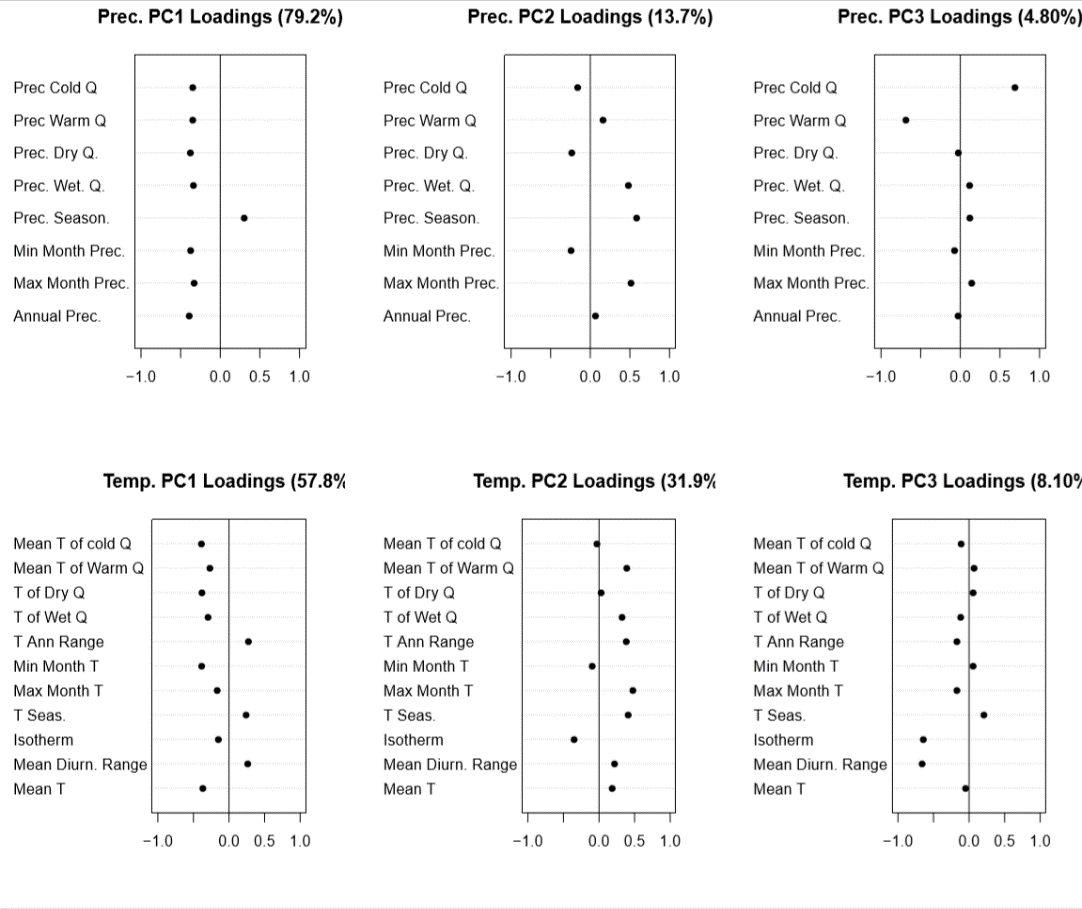

Figure S6: Principal component weights for precipitation and temperature. These are the principal components which were used for PGLS models. PC1 of Temperature explained 57.8% of variation in bioclim 1-11, and reflected temperature seasonality at high values and high mean temperature at low values. PC2 and PC3 of Temperature explained 31.9% and 8.10% of the variation and reflected variation along axes of mean and maximum quarterly temperatures and isothermality respectively. PC1 of Precipitation explained 79.2% of the variance in precipitation variables and mainly described variation between low and high mean seasonal precipitation. PCs 2 and 3 of Precipitation explained 13.7% and 4.80% of the variance in precipitation variables and best described variation in seasonality and precipitation in the coldest and warmest quarters respectively.
