## Supplementary figures and images for "Macroevolutionary bursts and constraints generate a rainbow in a clade of tropical birds"

### Supplementary Lorius Animation

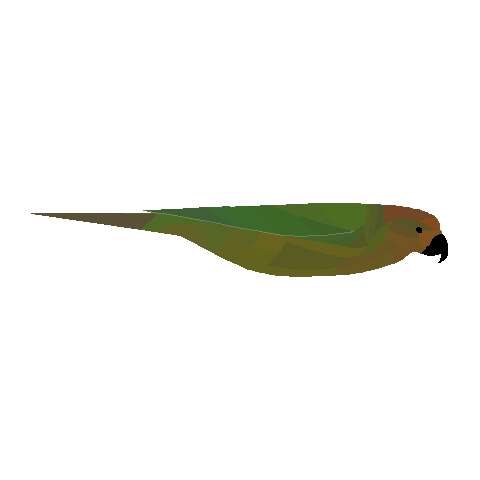
